# The first interneuron of the mouse visual system is tailored to the natural environment through morphology and electrical coupling

**DOI:** 10.1101/2024.06.12.598640

**Authors:** Matteo Spinelli, Alejandra Acevedo H., Christoph T. Block, Lucia Lindenthal, Fabian Schuhmann, Martin Greschner, Ulrike Janssen-Bienhold, Karin Dedek, Christian Puller

## Abstract

The topographic complexity of the mouse retina has long been underestimated, as obvious specializations, like a fovea or visual streak, are absent. However, anatomical and functional gradients exist. It was recently shown that receptive fields of retinal ganglion cells change their shape along the dorso-ventral retinal axis. These variations likely reflect the non-uniform statistics of the visual environment which vary dramatically from ground to sky. Horizontal cells are the first visual interneurons and dictate the synaptic signaling between photoreceptors and bipolar cells by lateral interactions, thereby shaping the receptive fields of down-stream neurons. Thus, we asked whether regional specializations are present at this earliest stage of synaptic circuitry, i.e. at the level of horizontal cells. We analyzed horizontal cell density distributions, morphological properties, localization of gap junction proteins, and the spatial extent of horizontal cell electrical coupling across complete retinas. All of these horizontal cell key features were asymmetrically organized along the dorso-ventral axis. Dorsal horizontal cells were less densely distributed, had larger dendritic trees, and electrical coupling was more extensive than in their ventral counterparts. The steepest change along this gradient occurred at the opsin transition zone of photoreceptors, i.e. the mouse visual horizon. Therefore, our results show that the cellular and synaptic organization of the mouse visual system are adapted to the visual environment at the earliest possible level, and that horizontal cells are well suited to form the cellular substrate for the global gradient previously described for the receptive field structures of retinal ganglion cells.

## Introduction

The topographic organization of neurons in the vertebrate retina reflects the lifestyle and behavior of a given species in its corresponding habitat. Some of the most basic functions of the visual system are supported by distinct cell distributions, such as prey capture or threat detection. One of the most prominent specializations is the human fovea, which provides us with high acuity vision. There, the cell density is extremely high, and the dendritic and corresponding receptive fields of the neurons are accordingly small. Our fovea is only one example of many, as topographic specializations can be found in various other vertebrate species (reviewed in Baden et al., 2020; Heukamp et al., 2020).

One of the most commonly used model systems for the early visual system in vertebrates is the mouse retina. There, a well-known topographic specialization is the pattern of opsin expression in cone photoreceptors (Röhlich et al., 1994; Applebury et al., 2000; Nadal-Nicolás et al., 2020). Green- and UV-sensitive opsins are expressed in a gradient along the dorso-ventral axis with a steep transition zone at the visual horizon of the mouse, reflecting the natural scene properties of its environment (Baden et al., 2013; Qiu et al., 2021). However, the mouse retina has long been thought to lack any further major topographic variations beyond the distribution of opsins. In line with this simplified anatomical perspective, and due to the lack of better knowledge, functional properties of a given cell type have also been assumed to be the same, or at least very similar, across different retinal regions.

Today it is well known that the mouse retina exhibits many different patterns of topographic variations, some are restricted to distinct cell types, whereas others are more global (reviewed in Heukamp et al., 2020). For instance, retinal ganglion cells show a distinct global density gradient with many more cells located in the ventral retina, i.e. the upper visual field (Salinas-Navarro et al., 2009; Duda et al., 2023). This is accompanied by complex density arrangements of cells in a type-dependent manner (e.g. Zhang et al., 2012; Bleckert et al., 2014; Rousso et al., 2016; Berry et al., 2023; Duda et al., 2023). Functional specializations of individual ganglion cells of a given type across different retinal locations go hand in hand with the anatomical ones and are equally complex (e.g. Chang et al., 2013; Sabbah et al., 2017; Warwick et al., 2018; Holmgren et al., 2021; Johnson et al., 2021). While some of these functional gradients may be shaped by intrinsic features of ganglion cells (Raghuram et al., 2019; Werginz et al., 2020), regional changes of their response properties typically originate in the presynaptic circuitry. However, topographic variations of cells upstream of retinal ganglion cells are poorly understood.

One of the most outstanding examples of a functional gradient in retinal ganglion cells is the dorso-ventral change of their receptive field surround structure in terms of amplitude and spatial extent, with remarkable asymmetries in the vicinity of the visual horizon (Gupta et al., 2023). The origin of this phenomenon remains unknown. Inhibitory amacrine cell signaling in the inner retina contributes to the formation of ganglion cell receptive field surrounds (reviewed in Diamond, 2017). Nevertheless, horizontal cells in the outer retina also play a fundamental role in the establishment of the ganglion cell receptive fields, including surround properties (Chaya et al., 2017; Drinnenberg et al., 2018; Ströh et al., 2018). Thus, we hypothesized that horizontal cells contribute to the region-dependent changes of ganglion cell receptive fields.

Horizontal cells are the first inhibitory interneurons in the vertebrate visual system and positioned to indirectly modulate ganglion cell response properties via lateral synaptic interactions with photoreceptors and bipolar cells in the outer retina (reviewed in Thoreson and Mangel, 2012). The spatial extent of lateral interactions is determined by horizontal cell dendritic tree size and extensive electrical coupling of dendrites by gap junctions (e.g. Bloomfield et al., 1995; Hombach et al., 2004; Shelley et al., 2006; Zhang et al., 2011). Our analysis of horizontal cell densities and dendritic field sizes, as well as gap junction distribution and the spatial extent of horizontal cell electrical coupling revealed a topographical organization of these features along the dorso-ventral axis of the retina. The most prominent distribution asymmetries occurred at the opsin transition zone, i.e. the mouse visual horizon.

## Material & Methods

### Animals and tissue preparation

All procedures were performed in accordance with the law on animal protection (*Tierschutzgesetz*) issued by the German Federal Government and approved by the local animal welfare committee. Mice of either sex were used, including the wild-type (C57BL/6J), Cx57^lacZ/lacZ^, and Cx57^+/+^ animals on C57BL/6J genetic background (Hombach et al., 2004). Animals were housed under standard conditions, including a 12-hr light/dark cycle with water and food *ad libitum*.

Mice (ages: 2-4 months for quantification of the horizontal cell (HC) density, 3-5 months for immunohistochemistry, 3-4 months for HC injections) were deeply anesthetized with carbon dioxide and killed by cervical dislocation or decapitation. Eyes were immediately enucleated and lens and vitreous were removed in a 0.01 M phosphate buffered saline (PBS, pH 7.4) or in a 0.1 M phosphate buffer (PB, pH 7.4). For HC injections, animals have been dark adapted for at least 1 hour before they were sacrificed and the subsequent tissue preparation was performed under infrared illumination using night vision goggles (No. G18597, Gutzeit-Gmbh). Retinal dissection was performed in Ames medium (Sigma/Biomol) supplemented with sodium bicarbonate and bubbled with carbogen at 30-32 °C, pH 7.4. Fixation of the tissues was performed at room temperature (RT). Posterior eyecups were immersion-fixed in initially cold 2-4% paraformaldehyde (PFA) diluted in PBS for 15-20 min. Injected retinas were fixed in 2% PFA diluted in 0.1 M PB for 20 min.

For immunohistochemistry, retinas were cryoprotected after fixation in sucrose solution (30% w/v) overnight at 4°C and then stored at -20°C until use. To keep track of the retinal orientation, the choroid fissures were used to place marking cuts into the tissue (Stabio et al., 2018). For cryosections, the tissue was then embedded in Tissue-Tek O.C.T. Compound (Sakura Finetek) and sectioned vertically at 20 µm using a Leica CM1860 cryostat. For whole-mounted retinas, four radial relieving cuts were made without compromising the initial marking. Then, the retinas were dissected in PBS or in Ames depending on the following set of experiments. This was performed in a way that the complete retina was preserved, including the most peripheral region (outer marginal zone). Finally, the tissue was mounted on a black nitrocellulose filter membrane (Millipore) with the ganglion cell side up, for immunohistochemistry or cell injections.

### Immunohistochemistry

Immunohistochemical labeling was performed by an indirect fluorescence method. Vertical sections were incubated overnight at RT with primary antibodies (Table 1 and supplemental information, Fig. S1) diluted in incubation solutions containing either 5% normal donkey serum (NDS), 1% bovine serum albumin (BSA), 0.5% Triton X-100 in PBS, or 5% Chemiblocker (Millipore) and 0.3% Triton X-100 in Tris-buffered saline (TBS, pH 7.6). Afterward, sections were incubated for 90 min with secondary antibodies diluted in the corresponding incubation solution. Whole-mounted retinas were incubated at RT for 2-3 days in the primary antibody solution containing 5% NDS, 1% BSA, and 1% Triton X-100 in PBS. Secondary donkey antibodies were incubated at RT for 2h or overnight in the same incubation solution. Type and dilution of secondary antibodies was the same for vertical sections and whole-mounts (Alexa Fluor 488, 568, and 647, 1:500, Invitrogen; Alexa Fluor 405, 1:500, Abcam; CF568, 1:500, Sigma).

**Table 1.** Primary antibodies.

| **Antibody** | **Host, type** | **Working dilution** | **Immunogen** | **Source, Catalog #** |
| --- | --- | --- | --- | --- |
| Calbindin | Rabbit, Polyclonal | 1:2000 | Recombinant rat calbindin D-28k | Swant CB-38A |
| Cx57 | Guinea pig, Polyclonal | 1:500(w)/1:100(c) | C-terminal peptide of mouse Cx57<br>PGSRKASFLSRLMSEK | This study |
| Cx57 | Guinea pig, Polyclonal | 1:200(w)/1:100(c) | C-terminal peptide of mouse Cx57<br>CSMSMILELSSIMKK | This study |
| Cx57 | Rabbit, Polyclonal | 1:500 | CSMSMILELSSIMKK | Janssen-Bienhold et al., 2009 |
| GluK1 | Mouse, Monoclonal | 1:2000 | aa 869-918 C-terminus of human GluK1 | Santa Cruz Biot. sc-393420 |
| S-opsin | Goat, Polyclonal | 1:10000 | N-terminus of the of human OPN1SW | Santa Cruz Biot. sc-14363 |
| ZO-1 | Mouse, Monoclonal | 1:100 | aa 334–634 of human recombinant ZO-1 | Zymed, 33-9100 |
| ZO-1 | Rabbit, Polyclonal | 1:100 | aa 463–1109 of human zonula occludens-1 cDNA | Zymed, 61-7300 |

Dilutions used for labeling of cryosections (c) or whole-mounted retinas (w) if different depending on condition

Retinas where HCs were injected with neurobiotin (see “Horizontal cell injections” below), which were intended for immunostaining of Cx57, were fixed and labeled using the same protocol as described above. Alexa Fluor 568-conjugated streptavidin (1:250, ThermoFisher Scientific) was used to visualize neurobiotin during the secondary antibody incubation.

Retinas were mounted together with the filter paper on slides in Vectashield (Vector Laboratories). Spacers between glass slides and coverslips were used to avoid squeezing the tissue. Dye-injected specimens were directly mounted on glass slides with Vectashield and coverslipped with spacers as described above. Neurobiotin-injected specimens for the tracer-coupling analysis of HCs were first incubated overnight at 4°C with Alexa568-conjugated streptavidin (1:250, Invitrogen, S11226), diluted in incubation buffer containing 10% NDS, 0.3% Triton X-100 in 0.1 M PB before mounting.

### Retina reconstructions

For the large-scale quantification of HC densities, the merged tile scans of image stacks from whole retinas were used. The R package Retistruct (Sterrat et al., 2013) was used to reconstruct the dissected whole-mounts to the spherical cap shape of the intact retina. 3D reconstruction allowed a better estimate especially at the cloverleaf outline and avoided some miscalculation introduced by stress during flattening. In each retina, the tissue outline and the incisions were manually marked. In rare cases of folded, damaged, or missing retinal tissue, the extent of the retina was manually estimated. The rim angle of the reconstructed retina was set to 110° and the radius of the retina to 1.5 mm, as measured from independent cryosections and in accordance with earlier reports (Schmucker and Schaeffel, 2004). The reconstructed spherical caps were visualized as an azimuthal equal-distance projection. Individual calbindin-positive HC bodies were manually marked using the CellCounter plugin in Fiji, in 5 retinas. The boundary of the short-wavelength sensitive (S-)opsin gradient was manually identified in the same 5 retinas and was represented as mean ± standard deviation (SD, Fig. 1, 5, S2, S3). The local density of horizontal cells was calculated in spherical coordinates by counting the number of cells in a 10 degree spherical cap. This radius corresponds to an arc length of ∼260 µm and a counting disc area of 0.21 mm^2^. This counting area was corrected to reflect marked regions where data was unattainable. The average density across different retinas was calculated in a fixed regular grid.

**Figure 1.**
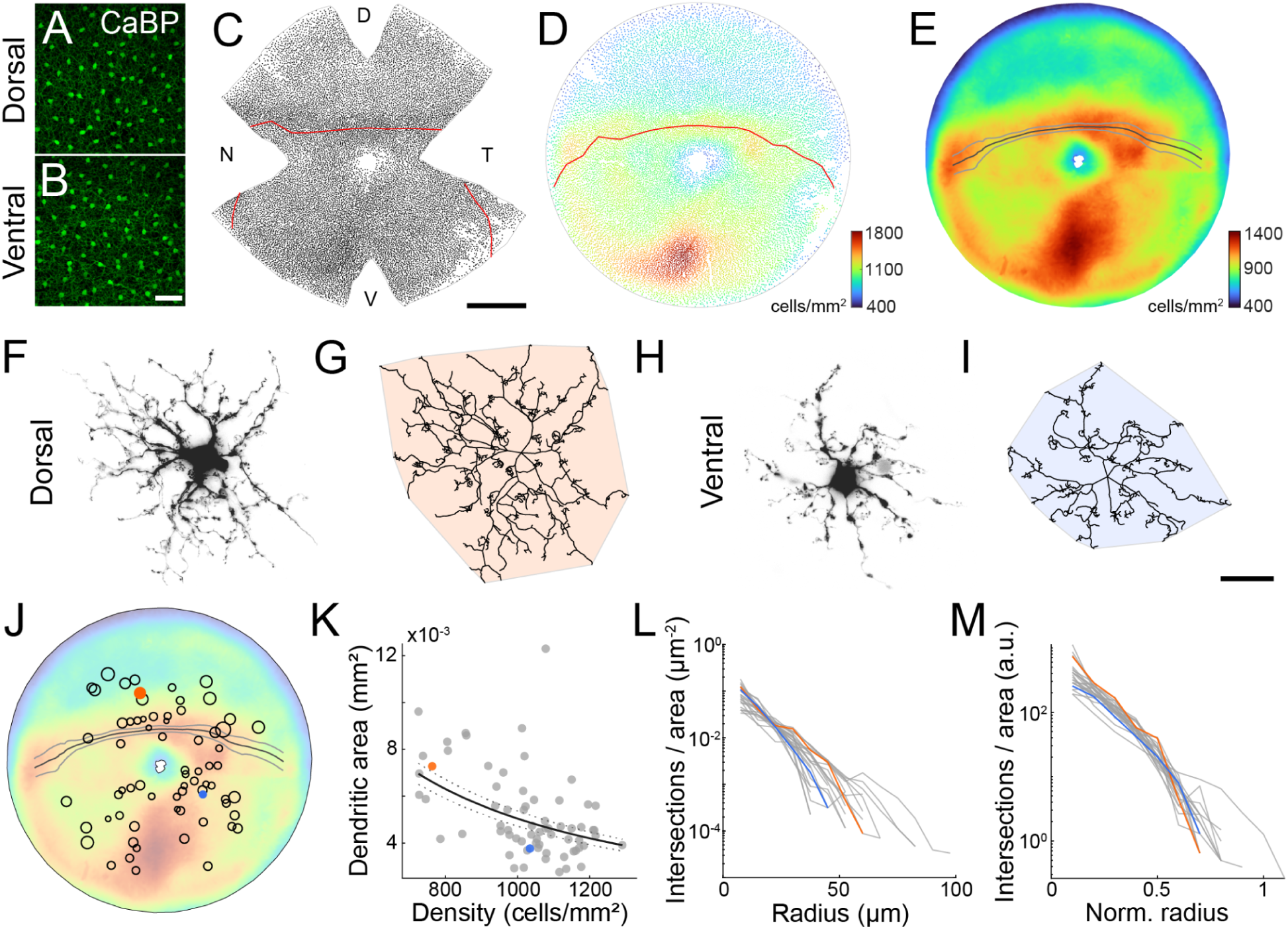
Horizontal cell density and dendritic field size change across the retina. **A,B:** Confocal images of dorsal (A) and ventral (B) whole-mounted retina labeled against calbindin (CaBP). **C:** Each dot represents the cell body position of a horizontal cell marked with CaBP across a complete retina of a left eye. The red lines in C and D mark the S-opsin transition zone (see also Fig. 4). D, dorsal; V, ventral; N, nasal: T, temporal. **D:** Azimuthal equal-distance projection of the reconstructed retinal sphere of the data shown in C. The color of the dots represents the local density of the corresponding cell body position. **E**: Horizontal cell density distribution averaged across 5 retinas. Black line, mean S-opsin transition zone, gray lines, SD (n=5). Color bars in D and E indicate cells/mm^2^. **F-I:** Dye-injected horizontal cells from dorsal (F) and ventral (H) retina, together with the corresponding skeletons (G, I) traced through confocal image stacks of the cell with convex hulls to determine the dendritic tree area. **J:** Positions and dendritic field sizes of 69 injected horizontal cells, superimposed on the averaged density distribution from E. Circle area represents the relative area of the corresponding horizontal cell; for better visibility circles are not shown to scale. Filled orange and blue circles show the positions of horizontal cells from F and H, respectively. **K:** Relation between horizontal cell dendritic field size and local horizontal cell density. Lines describe the best estimate assuming a constant coverage factor (5.04, 95% CI [4.72, 5.37]). **L:** Sholl analysis of dendritic intersections from 23 horizontal cells. **M:** As in L but data normalized to dendritic area (convex hull). Scale bars: 50 μm in B, 1 mm in C, 20 μm in I, applies to F-I.

**Figure 2.**
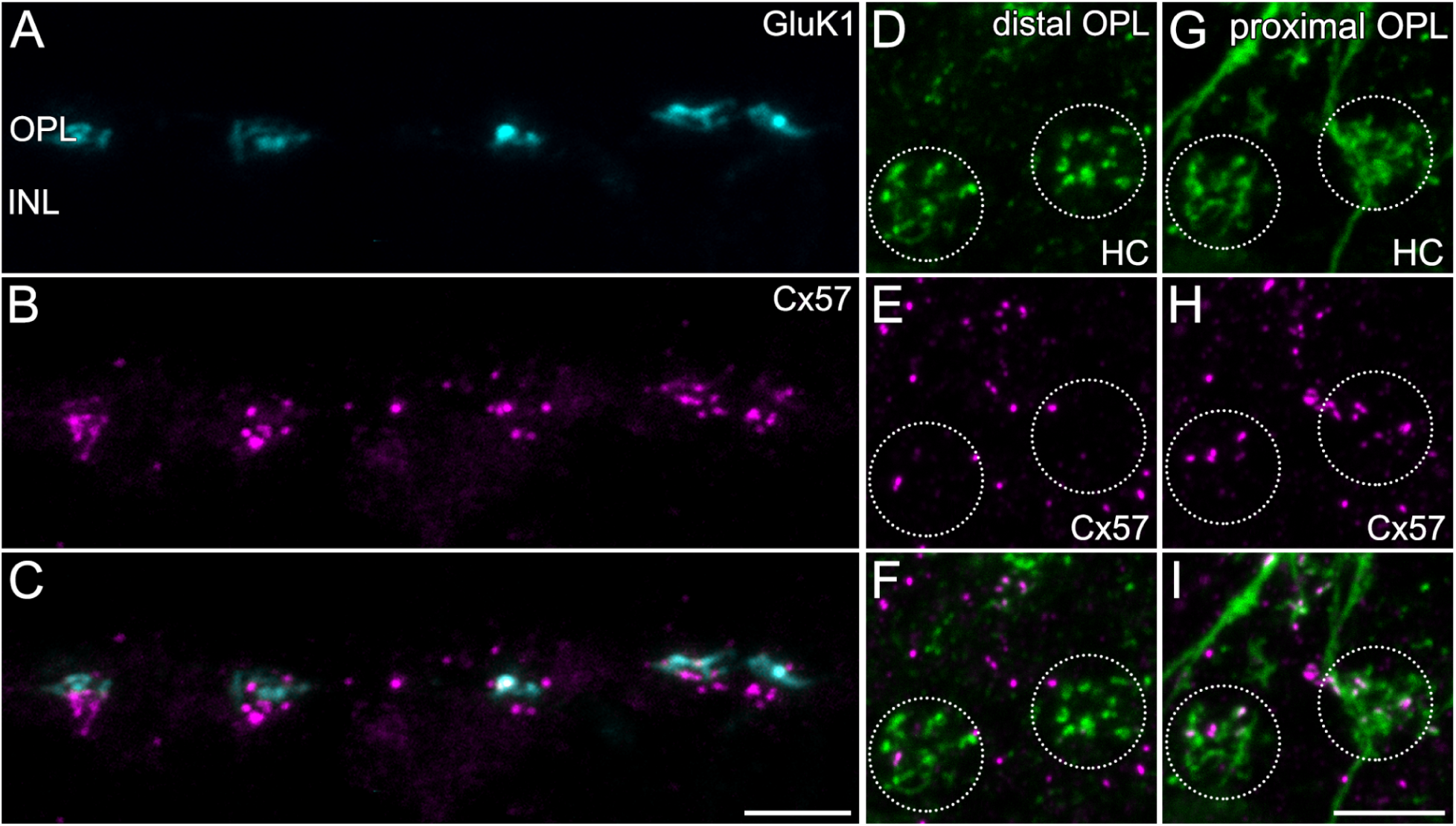
Spatial localization of horizontal cell gap junctions. **A-C**: Projections of a confocal image stack of a vertical cryostat section double-labeled for the kainate receptor subunit 1 (GluK1, used as a cone pedicle marker) and the horizontal cell gap junction protein connexin 57 (Cx57). **D-F:** Maximum intensity projections of a confocal image stack of whole-mounted retina labeled with antibodies against Cx57 and with Alexa Fluor 568-conjugated streptavidin after microinjection of neurobiotin into a horizontal cell. The distribution of Cx57 is shown at the tips of horizontal cell (HC) dendrites invaginating into two cone pedicles. Pedicle positions were identified by these clusters of invaginating dendritic tips and are indicated by dashed circles. **G-I:** As in D-F, but ∼2 μm beneath the same pedicle position. INL, inner nuclear layer; OPL, outer plexiform layer. Scale bars: 10 μm in C, applies to A-C; 5 μm in I, applies to D-I.

**Figure 3.**
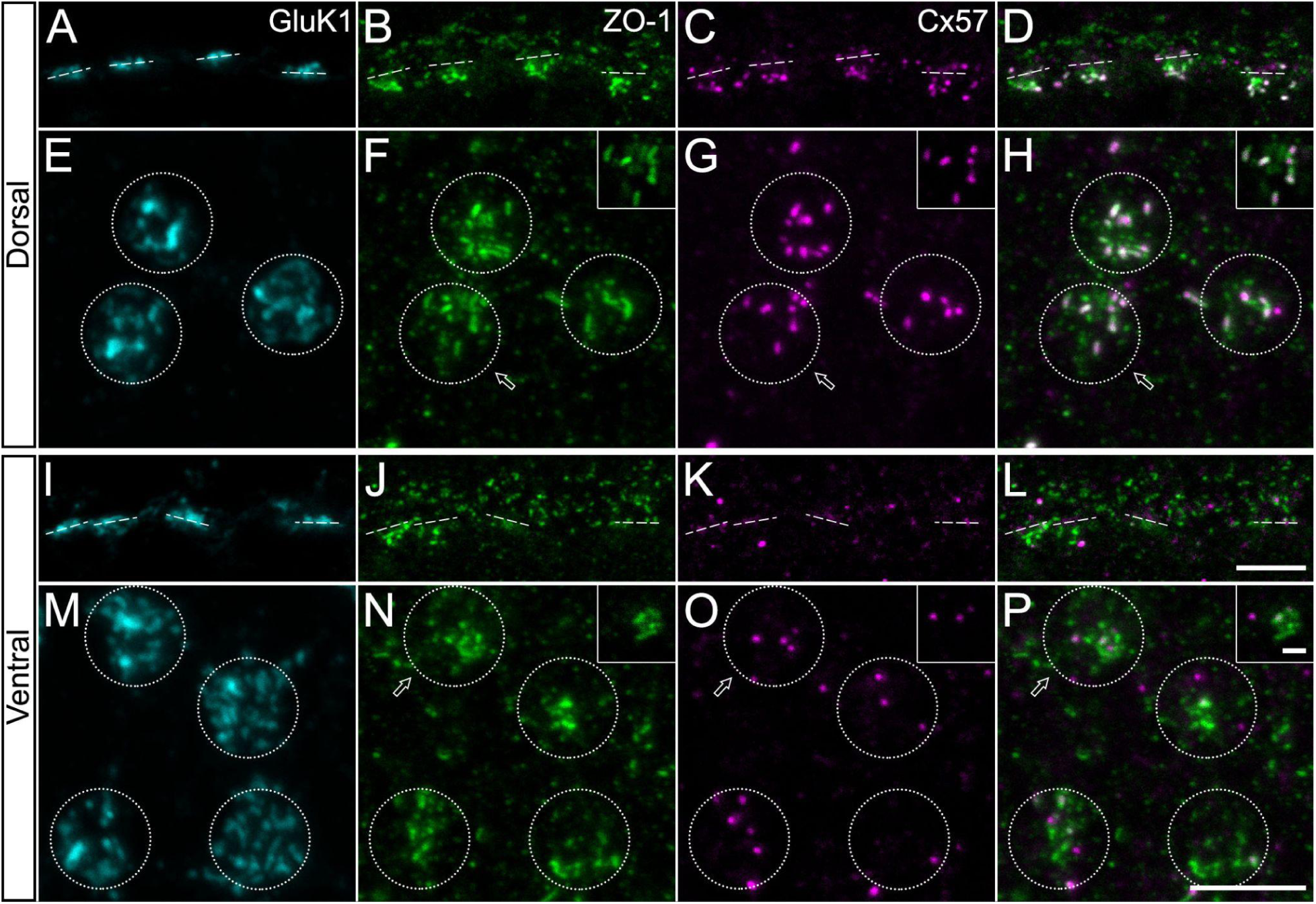
Clustering of ZO-1 and Cx57 differs in dorsal and ventral retina. **A-D:** Maximum intensity projections of a confocal stack from a vertical cryostat section of dorsal retina triple-labeled against the kainate receptor subunit 1 (GluK1), the tight junction protein zonula occludens-1 (ZO-1), and connexin 57 (Cx57). Dashed lines indicate pedicle positions marked by GluK1 staining. **E-H:** As in A-D from a piece of whole-mounted retina. Maximum-intensity projections were chosen to cover the proximal OPL beneath cone pedicles. Circles indicate pedicle positions. Arrows indicate an example pedicle of which single optical sections of the corresponding channels are shown in the box (top right). **I-P:** As in A-H but from ventral retina. Scale bars: 5 μm in L, applies to A-D and I-L; 5 μm in P, applies to E-H and M-P; 1 μm box in P, applies for single optical section boxes E-H and M-P.

Coverage factor was defined as the number of HCs within the dendritic field of one cell and estimated by multiplication of density and convex hull area. Under the assumption of a constant coverage factor, this relation was fitted in MATLAB using the data shown in Fig. 1K.

### Horizontal cell injections

Cell nuclei in whole-mounted retinas were visualized by prior incubation in Ames medium containing 0.3 - 0.5 μM of DAPI (Abcam) for 60 min at 32 °C. Then the retinas were mounted onto black nitrocellulose membrane (see above).

Borosilicate glass electrodes were pulled with a micropipette puller (P-97, Sutter Instrument CO) to obtain sharp electrodes with a resistance between 100 – 200 MΩ. HCs were injected either with a fluorescent dye alone (to analyze the morphology of the cells) or with a mixture of dye and the gap junction tracer molecule neurobiotin (to analyze the electrical coupling).

For fluorescent dye injections, the electrodes were filled with 2 µl of 5 mM of Alexa Fluor 568 hydrazide (Invitrogen, diluted in 200 mM KCl) and backfilled with 10 µl of 200 mM KCl. Epifluorescence light was used to identify DAPI-labeled HC bodies based on their location and their relatively large size. Candidate cell bodies were targeted under visual control with epifluorescence illumination and impaled with sharp electrodes for dye iontophoresis using -0.5 nA square pulses of 500 ms at 1 Hz for 3 min. After injections, the retina was fixed in PFA as described above.

Neurobiotin injections were restricted to the nasal side of a given retina. Electrodes were tip-filled with 3 µl of a 1:1 mixture of 4% neurobiotin (SP1120, Biozol) diluted in 0.1 M Tris buffer (pH 7.3) and 5 mM Alexa Fluor 568 Hydrazide, and back-filled with 10 µl of 200 mM KCl in Tris buffer. The dye was injected as described above for 1-2 minutes before the current was reversed to inject neurobiotin using +0.5 nA square pulses of 500 ms at 1 Hz for 5 min. Then, a neurobiotin diffusion time of 10 min was granted before fixation. Only a single HC was injected per retina (dorsal n=7, ventral n=7) to provide constant conditions for these experiments to quantify tracer coupling.

### Image acquisition

High-resolution fluorescence image stacks were acquired using TC SP8 or TCS SL confocal laser scanning microscopes (Leica). Scanning was performed with 63x/1.32 or with 63x/1.4 oil-immersion objectives and z-axis increments between 0.1 and 0.3 µm. The SP8 confocal microscope with either 40x/1.3 or 63x/1.4 oil-immersion objectives was also used to acquire image stacks of HC dendritic trees. Image stacks of the coupled HCs were acquired using a 20x/0.70 oil-immersion objective.

Images of entire whole-mounted retinas were obtained with a Leica DM6B epifluorescence microscope equipped with a motorized stage and a 20x/0.5 air objective. Individual image stacks of a tile scan were automatically stitched together in the microscope software (LAS X, Leica). Overviews of the entire retinas were used for the large-scale quantification of HC density and to keep track of the precise location of the injected horizontal cells (see “retinal reconstruction” section).

Images are presented as single optical sections or as maximum intensity projections of image stacks. Some images were further processed with Fiji (Schindelin et al., 2012) using the “subtract background” (rolling ball) plugin and intensities were normalized using the “enhance contrast” plugin with 0.01% saturation. Brightness and contrast of the final images were adjusted using Photoshop (Adobe).

### Tracing and morphometric analysis

Tracer-coupled HCs were manually counted in Fiji using maximum intensity projections of confocal image stacks. Area measurements were based on convex hulls either encompassing dendritic trees of the HCs (Fig. 1) or all neurobiotin-positive cell bodies (Fig. 5).

Horizontal cell coupling was studied in 14 retinas and the positions and coupling strength were shown along with the independently measured horizontal cell density. The dendritic area was studied in 69 injected cells from 20 retinas (18 animals, either eye). The area was measured from convex hulls of the manually marked dendritic tree. A Sholl analysis (Sholl, 1953) was used to compare the dendritic branching pattern of traced HCs (n=23) across retinal regions. For this, HC skeletons were traced through image stacks using Amira (Thermo Scientific). The number of intersections of dendritic processes with concentric circles was divided by the area of the respective circle. In Fig. 1M, each cell was normalized to equal convex hull area to focus only on the arborization patterns, not on sizes. Cells with overlapping, co-injected cells or any interference from blood vessels, which hindered accurate interpretation, were excluded.

### Colocalization analysis

Immunoreactivity of Cx57 and ZO-1 was measured to assess spatial extent and colocalization. Confocal stacks of cone pedicles from the dorsal and ventral periphery of three retinas were analyzed. The total extent in z direction was chosen to capture the entire thickness of the ZO-1 cluster beneath a given cone pedicle (Puller et al., 2009), which typically included 20-25 consecutive optical sections. For each cone pedicle, a circular region of interest (ROI) with a diameter of 7 µm was selected around the ZO-1 cluster. Background and contrast was adjusted as described above, and a global threshold was independently applied. Colocalization of Cx57 and ZO-1 was analyzed with the “colocalization highlighter” plugin in Fiji (MBF collection, Collins, 2007; Tetenborg et al., 2017). Areas of colocalization and areas of individual immunostaining per channel were measured in the same ROIs with the “analyze particles” plugin in Fiji. Colocalization areas smaller than 0.01 µm^2^ were excluded from the analysis. Colocalization in images with one vertically flipped channel per ROI at a given cone pedicle position served as control measurements (Puller et al., 2007). For the quantification of the individual immunostainings, particles with a size smaller than 0.04 µm^2^ were excluded from the analysis. This analysis was applied to 3 retinas. The area measurements were normalized (Fig. 4E-G) to account for differences between the samples regarding staining intensity and the relative background staining levels.

**Figure 4.**
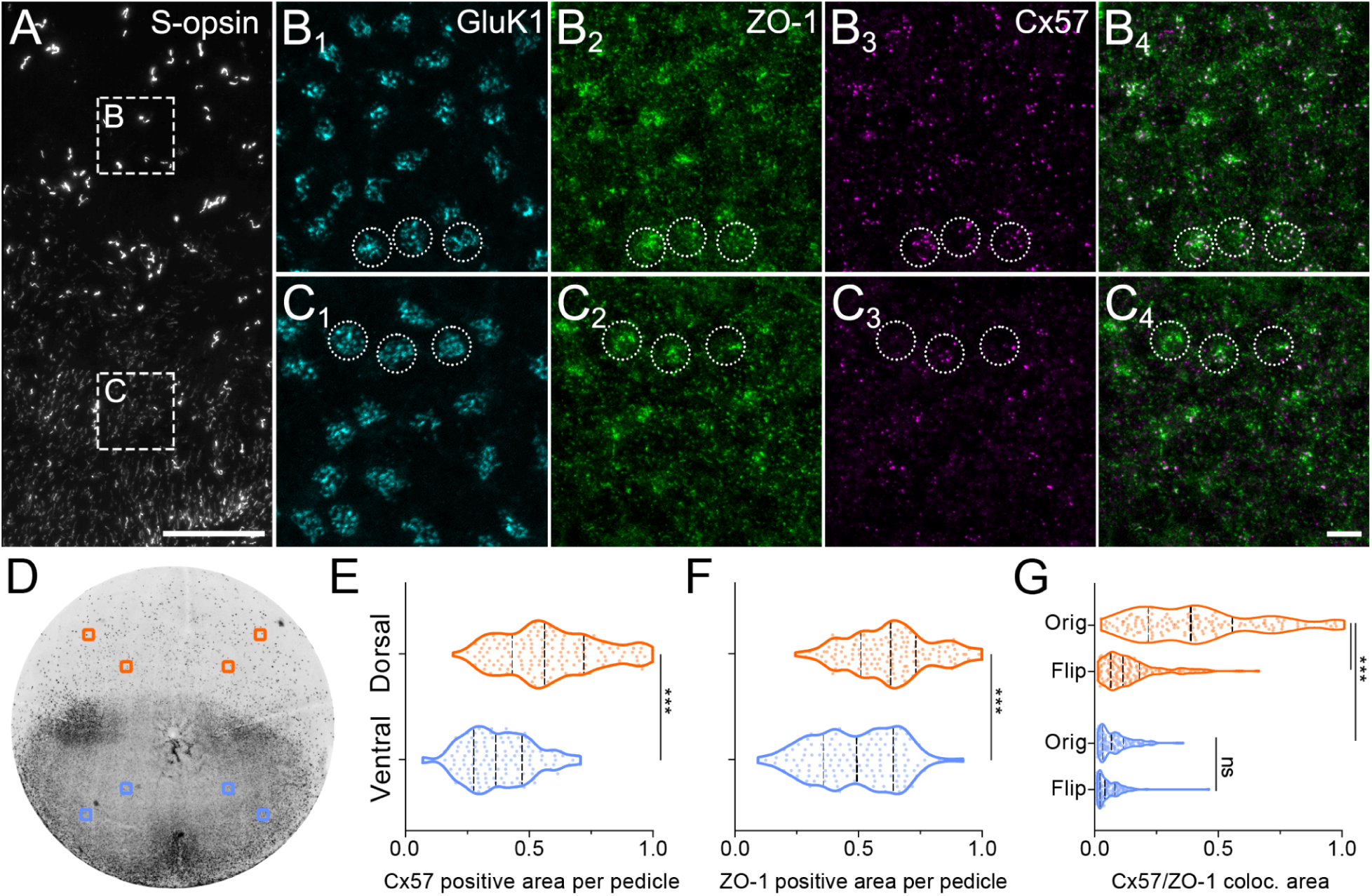
Clustering density of horizontal cell gap junctions changes at the S-opsin transition zone. **A-C:** Maximum intensity projections of confocal image stacks from the region of the short-wavelength sensitive (S)-opsin transition zone. The retinal whole-mount was quadruple-labeled with antibodies against S-opsin, GluK1, ZO-1, and Cx57. A, cone outer segments positive for S-opsin at the transition zone. Boxes B and C indicate regions where confocal image stacks (B_1_-C_4_) were acquired. B_1_-B_4_ shows GluK1, ZO-1, and Cx57 at the level of the OPL in the position of box B (in A), close to the transition zone but on its dorsal side. Three example pedicles are indicated by circles. C_1_-C_4_, as in B_1_-B_4_ but for the area C on the ventral side of the transition zone. **D:** Azimuthal equal-distance projection of tile scans of a complete, whole-mounted retina immunolabeled against S-opsin. Boxes indicate the locations where image stacks were acquired for the quantification of Cx57 and ZO-1 immunostaining. **E-G:** Quantification of the immunolabeled areas (normalized from µm^2^) of Cx57 and ZO-1 and their colocalization per circular ROI beneath each pedicle. Measurements were pooled across dorso-peripheral locations (orange, all upper boxes in D combined) and the ventro-peripheral locations (blue, all lower boxes in D combined). Dashed lines in violin plots show median and quartiles. Each data point represents the measurement in a ROI beneath one pedicle. Orig, colocalization analyzed in the original image stack, Flip, control measurement where one channel per ROI was vertically flipped. *** p<0.001; ns, not significant. Scale bars: 100 μm in A; 5 μm in C_4_ applies to B_1_-C_4_.

### Statistical analysis

Statistical tests were performed using Prism 9 (GraphPad Software) on normalized ZO-1 and Cx57 immunostaining data (Fig. 4). A one-way ANOVA Tukey for multiple comparisons was used to compare the colocalized area in the dorsal and ventral periphery. Flip controls were used to account for randomly observed colocalization. An unpaired, two-tailed Mann Whitney test was used to compare the immunoreactive areas of Cx57 and ZO-1 between dorsal peripheral and ventral peripheral measurements. A p-value < 0.05 was considered statistically significant. Quantitative data was obtained from 3 retinas including 144 dorsal pedicles, 131 ventral pedicles presented as median and quartiles.

A Mann-Whitney test (Prism 7, GraphPad Software) was performed to test for statistical significance (p<0.05) between the number of coupled cell bodies and the area covered by them (Fig. 5). Quantitative data was obtained from 14 retinas (7 retinas for each dorsal and ventral sides) reported as mean ± SD in the text description and the median is indicated in Fig. 5D, E.

**Figure 5.**
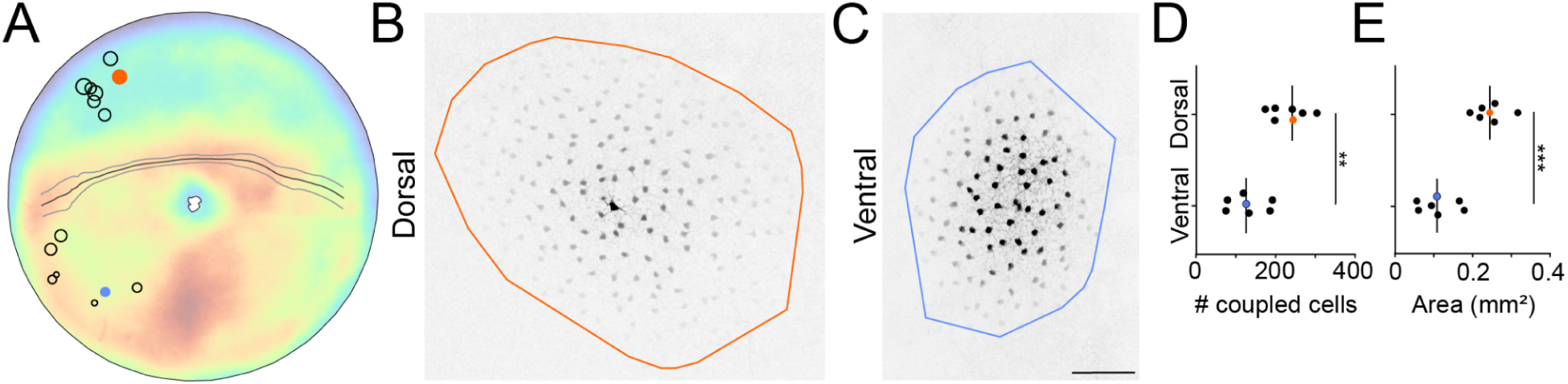
Tracer coupling patterns of horizontal cells differ in dorsal and ventral retina. **A:** The position of neurobiotin-injected horizontal cells is indicated by circles superimposed on the density distribution disc taken from Fig. 1E. The area of the circles reflects the number of tracer-coupled cell bodies per injected horizontal cell. Orange and blue filled circles represent the example injections in B-E. **B,C:** Example tracer-coupling patterns upon injection of neurobiotin into single cell bodies (B, in dorsal retina; C, ventral). Convex hulls encompassing all neurobiotin-positive cell bodies were used to measure the area of neurobiotin-spread. **D,E:** Quantification of the number of tracer-coupled horizontal cell bodies (D, p<0.01) and the area of tracer-spread (E, p<0.001) per injection. Vertical lines indicate median. Scale bar 100 μm in C, applies to B and C.

## Results

Horizontal cells play a crucial role in shaping the retinal output signals (Chaya et al., 2017; Drinnenberg et al., 2018; Ströh et al., 2018). Moreover, horizontal cells may match the ganglion cell receptive field surround properties with the visual environment: the strength and spatial extent increase along the dorso-ventral axis of the retina (Gupta et al., 2023). This change should then be reflected in horizontal cell morphological properties to support signaling at different spatial scales, i.e. large in dorsal retina, small in ventral retina, with a distinct transition zone at the visual horizon. Indeed, some evidence suggests that horizontal cells show a certain level of density variation across retinal regions (Camerino et al., 2021) which may affect the sizes of cells. Thus, we analyzed the horizontal cell density distribution in high detail by considering each and every cell body across complete mouse retinas.

### Horizontal cell density and dendritic tree size form a gradient across the retina

Whole-mounted retinas were labeled with antibodies against calbindin (CaBP, Fig. 1A, B), a common mouse horizontal cell marker (Haverkamp and Wässle, 2000). The positions of all immunolabeled horizontal cell bodies were manually marked in a merged microscopic tile scan of the flattened retina (Fig. 1C), where each dot indicates the position of a horizontal cell body (17650±1412, n=5 retinas, see supplemental Fig. S2). The cloverleaf shape of the tissue was reconstructed back into the original, almost hemispherical structure of the eyecup with Retistruct to calculate the densities of cells across the complete retinas (Sterratt et al., 2013; Duda et al., 2023). The local density was calculated and shown in an azimuthal equal distance projection (Fig. 1D). The 3D reconstruction allowed a proper averaging of data across five retinas independent of the relief cuts in the original whole-mounts. The retinas were counterstained with antibodies against the short-wavelength sensitive (S-)opsin to reveal the transition zone of opsin expression as an indicator of the visual horizon on the retina (Nadal-Nicolás et al., 2020; Qiu et al., 2021; Gupta et al., 2023). The transition zone is represented by lines in Fig. 1C-E (see Fig. 4A and D for representative microscopic images). Note that the transition zone was located far above the optic disc, but then it crossed the peripheral ends of the lower leaflets in a typical retinal whole-mount preparation (Fig. 1C). Future studies of retinal whole-mounts may be informed by these regionalization details and ensure a proper interpretation of data collected from definite positions of the mouse retina.

The horizontal cell density was low in the dorsal retina and high in the ventral retina. The steepest change in this gradient occurred in the area of the S-opsin transition zone. The general pattern of this density distribution closely resembled the density gradient of mouse ganglion cells (Salinas-Navarro et al., 2009; Duda et al., 2023).

Like other retinal cell types, horizontal cells form a mosaic across the retina, where the dendritic trees of the cells exhibit a constant overlap (Wässle and Riemann, 1978; Reese et al., 2005; Keeley et al., 2020). Thus, a change in cell density should yield a corresponding change in dendritic field sizes. Individual horizontal cells were dye-injected (n=69) to investigate if the density distribution serves as a direct read-out of dendritic field size, and to gain further insights into the horizontal cell dendritic field structure across the retina (Fig. 1F-J, supplemental Fig. S3). Convex hulls that encompassed the dendritic trees of the injected cells were used to measure their dendritic areas. The analysis revealed that horizontal cell dendritic trees located above the transition zone covered a much larger area (n=25, 6165±2357 µm^2^) than cells below (n=44, 4633±1196 µm^2^), with calculated dendritic field diameters of 87±2 µm or 76±9 µm, respectively (tailed Wilcoxon rank sum p<0.01). A fit to explain the relationship between the cell density and the dendritic area (Fig. 1K) predicted a coverage factor of ∼5, which is largely in line with previously published results (Reese et al., 2005). Horizontal cell morphology is characterized by dense dendritic branching in central parts of the dendritic tree and sparser, less arborized dendrites in the periphery (Raven et al., 2007; Behrens et al., 2022). Here, injected cells were traced through the image stacks (n=23, supplemental Fig. S3). A Sholl analysis (Sholl, 1953) was applied to the resulting skeletons to investigate whether this branching pattern of horizontal cells may differ between cells from different locations (Fig.1 L,M). This was not the case, as the results showed a constant branching pattern across the retina. The homogeneous morphology of horizontal cells across all dorsal and ventral cells in our sample became even more prominent (except for one outlier) when the data was normalized to account for the different sizes of the cells.

### Horizontal cell dendritic gap junction density changes at the visual horizon

The shape and size of a dendritic tree of a given retinal neuron is typically one of the major determinants of its functional receptive field. Thus, based on our findings, one would expect to find much larger horizontal cell receptive fields in the dorsal retina relative to those located below the opsin transition zone. On the other hand, horizontal cells are well known to exhibit extensive electrically coupled networks via gap junctions, which can lead to an increase in the spatial extent of receptive fields and lateral signal spread (Shelley et al., 2006; Zhang et al., 2011). Therefore, we aimed to analyze potential differences of horizontal cell dendritic gap junctions across the retina, which may further influence horizontal cell signaling properties in a region-dependent manner.

A prerequisite for such an analysis is, of course, a detailed knowledge of the types and positions of horizontal cell gap junctions. Electrical coupling of mouse horizontal cell dendrites is exclusively supported by gap junctions containing connexin 57 (Cx57; (Hombach et al., 2004; Shelley et al., 2006; Janssen-Bienhold et al., 2009; Puller et al., 2009). Together with Cx50, Cx57 is also involved in the formation of gap junctions between horizontal cell axon terminals (Dorgau et al., 2015). However, the latter are only connected to rod photoreceptor terminals and do not contribute to dendritic signaling (Trümpler et al., 2008). Therefore, axon terminal connexins were largely excluded from our study by restricting the analysis to mostly dendritic Cx57 in a volume beneath a given cone pedicle base (Fig. 2).

Individual dendrites of multiple horizontal cells converge beneath the cone pedicles in the proximal part of the outer plexiform layer (OPL) before their tips invaginate into the pedicle at the glutamate release sites. Mammalian horizontal cells are thought to form dendritic gap junctions primarily in this area of the proximal OPL (Puller et al., 2009). Correspondingly, Cx57-immunoreactive plaques were mostly clustered in the proximal OPL beneath the array of cone pedicles (Fig. 2A-C).

Horizontal cells were injected with neurobiotin, a small tracer molecule which can pass through gap junctions between cells to reveal their complete electrically coupled network (Vaney, 1991; Bloomfield et al., 1995; He et al., 2000; Zhang et al., 2011). Here, we combined neurobiotin-injected cells with immunolabeling of Cx57 and high-resolution fluorescence imaging to reveal the exact position of horizontal cell gap junctions on their dendrites (Fig. 2D-I). Cx57-positive plaques were colocalized with horizontal cell dendrites in a region ∼2 μm beneath the pedicle, where the dendrites converged and exhibited slight swellings. This may correspond to a specialized area where glutamate-receptor containing desmosome-like junctions are closely associated with gap junctions and the tight-junction protein zonula occludens-1 (ZO-1) on horizontal cell dendrites (Haverkamp et al., 2000; Puller et al., 2009).

Counterstaining of Cx57 with ZO-1 in the dorsal retina (Fig. 3A-H) confirmed previous results where the immunolabeling was largely colocalized beneath mouse cone pedicles (Puller et al., 2009). The same experiment performed on the ventral retina yielded a completely different picture (Fig. 3I-P). While the overall extent of GluK1 as a pedicle marker did not obviously change across regions, ZO-1 was less densely clustered and lacked the typical elongated plaques beneath pedicles and in areas between them. Most importantly, the amount of Cx57 immunoreactive puncta appeared largely reduced, resulting in a decrease of colocalization between Cx57 and ZO-1. Next, the distribution pattern of ZO-1 and Cx57 was investigated in detail relative to the S-opsin gradient with its transition zone as an indicator for the visual horizon of the mouse. It became obvious that the change from dense Cx57 clustering and robust colocalization with ZO-1 toward highly reduced Cx57 and the lack of colocalization occurred only within a few hundred microns at the transition zone (Fig. 4A-C). Colocalization of Cx57 with ZO-1 and measurements of the area covered by the corresponding immunostaining was performed in small image stack volumes collected from circular ROIs beneath each pedicle (“proximal OPL” as in Fig. 2G-I).

The data was collected and pooled from four locations in the dorsal periphery or in the ventral periphery, respectively (Fig. 4D; 3 retinas, 144 dorsal pedicles, 131 ventral pedicles). The original notion that ZO-1 and specifically Cx57 were less densely clustered at ventral cone pedicles was confirmed by the quantification of immunoreactive areas. These areas were measured within ROIs beneath a given pedicle. Thresholding and normalization was applied (Fig. 4E-G, see methods section for details) to account for different signal-to-noise ratios of the immunostainings in the 3 retinas. ZO-1 and Cx57-positive areas were reduced from dorsal to ventral retina by 22% and 35%, respectively (Fig. 4E,F). Furthermore, the colocalization of ZO-1 and Cx57 was reduced by 84% from dorsal to ventral parts of the retina (Fig. 4G).

As control measurements, colocalization was analyzed again in the same images but with one vertically flipped channel per ROI at a given cone pedicle position (e.g. Puller et al., 2007). Colocalization in the flip control of the dorsal retina was significantly reduced by 73%, suggesting that the overlap measured in the original image did not occur randomly.

Colocalization in control measurements of the ventral retina was reduced by 45%. This reduction was not significant due to the low amount of colocalization in the original images.

### Changes in gap junction organization translates into distinct patterns of electrical coupling

The difference in the gap junction density on horizontal cell dendrites from dorsal and ventral retina was striking. It remained unclear, however, whether this change resulted in a functional difference, i.e. different electrical coupling strengths in dorsal versus ventral regions. Therefore, horizontal cells were injected with the tracer molecule neurobiotin to visualize the extent of electrical coupling of a given cell in a determined location of dorso- or ventro-nasal retina (Fig. 5A). Beyond the spatial restriction to the nasal retina, each horizontal cell was injected under constant conditions, including current, injection duration, time of day, lighting, and temperature, to allow for proper comparability between the experiments. The results from these experiments confirmed distinct coupling patterns as suggested by our anatomical results. In the dorsal retina, more cells were electrically coupled to a given horizontal cell (n=7; 233±45 coupled cells) and the coupling extended across a larger area (0.245±0.039 mm^2^; Fig. 5B-E). The exact opposite was true for ventral horizontal cells (n=7), which exhibited coupling to fewer cells (130±46) in a smaller area (0.111±0.047 mm^2^).

## Discussion

Our study provides evidence for dorso-ventral asymmetries of horizontal cell features in the mouse retina, in terms of I) the cell density and dendritic tree size, II) the density of gap junctions clustered on their dendrites, and III) the extent of electrical coupling regarding the number of coupled cells and the area covered by them. Cell density gradients across the mammalian retina are well known, including corresponding morphological changes of the cells (Baden et al., 2020; Heukamp et al., 2020). It is remarkable, however, that the horizontal cell density gradient reported here is accompanied by a matching change in the synaptic architecture of the cells, i.e. the density and distribution of electrical synapses and the corresponding coupling strength. This change will directly affect the spatial extent of signaling at the first synapse in the visual system. Moreover, we provide evidence that these asymmetries match the global properties of the visual environment of the animal, where the most prominent transition of the aforementioned horizontal cell features occurs in the area of the visual horizon of the animal.

### Horizontal cells support the functional separation of the mouse visual field

Differences in the functional separation of upper and lower mouse visual fields are well known and some neuronal adjustments of the outer retina have already been revealed at the level of photoreceptors and bipolar cells (Röhlich et al., 1994; Applebury et al., 2000; Baden et al., 2013; Nadal-Nicolás et al., 2020; Camerino et al., 2021; Qiu et al., 2021; Sharpe et al., 2022). To our knowledge, horizontal cells have rarely been considered in this context (but see (Camerino et al., 2021, their Fig. 3). Horizontal cells located above the photoreceptor transition zone receive visual information from the ground, and they possess large dendritic trees and exhibit extensive electrical coupling. Horizontal cells below this zone are positioned for information from the upper visual field by smaller dendritic trees and reduced spatial extent of electrical coupling. Therefore, the characteristics of large dendritic trees paired with extensive coupling versus small dendritic trees paired with reduced coupling operate hand in hand as complementary phenomena to create large or small functional units in the lower or upper visual field, respectively (Fig. 6). Small functional units can serve high spatial resolution and would be beneficial for the detection of threats from overhead predators, for instance. Previous work has identified the transient OFF alpha ganglion cell as one of the retinal output cell types responsible for triggering innate defensive behaviors upon approach detection (Münch et al., 2009; Kim et al., 2020; Wang et al., 2021). These ganglion cells are prominently affected by horizontal cell signaling (Ströh et al., 2018), which is in line with the idea of horizontal cells contributing directly to the processing of the visual scene and to retinal output signaling. On the other hand, integration of signals across a larger area, specifically in the receptive field surround, may improve signal detection in front of the animal and in the lower visual field assuming more abundant transitions of low and high contrast signals.

**Figure 6.**
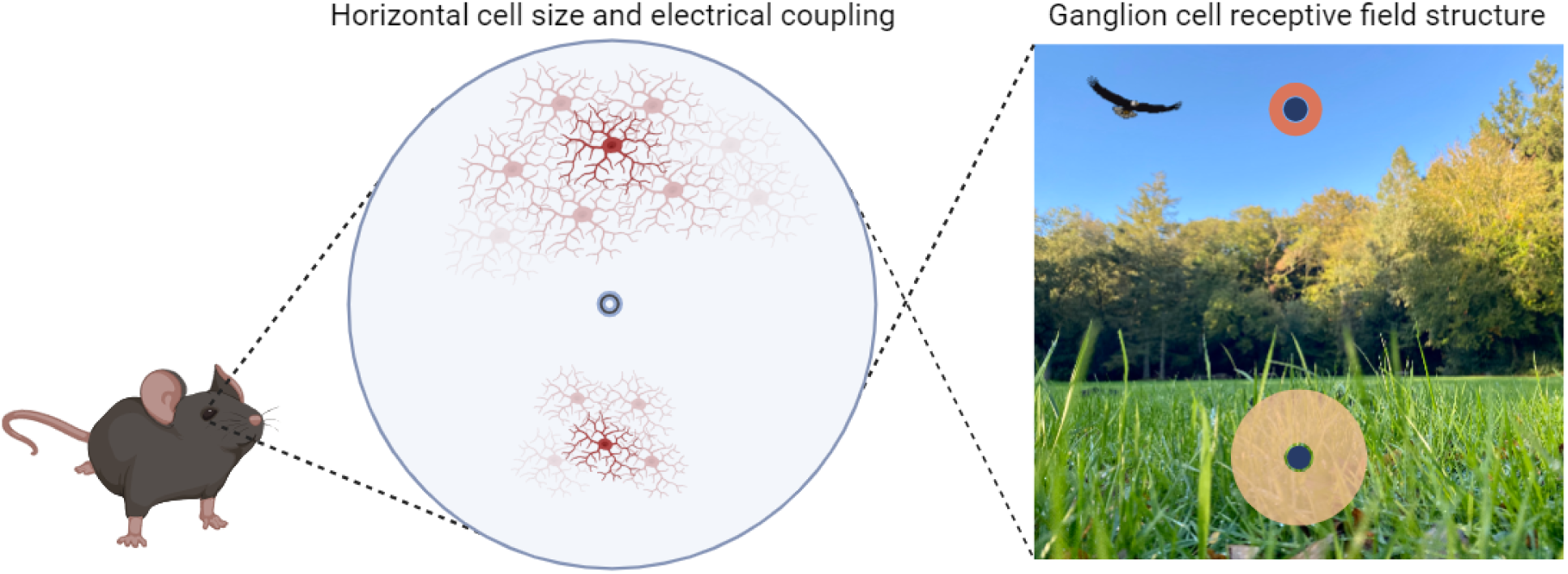
Horizontal cell feature asymmetry supports a receptive field architecture adapted to the visual environment. Dendritic tree size and electrical coupling area of horizontal cells is roughly twice as large in the lower visual field than in the upper. The feature asymmetries shown here are well suited to form the cellular substrate underlying the global asymmetry of ganglion cell receptive field surrounds across the retina (right, brown discs, blue discs represent receptive field centers, adopted from Gupta et al., 2023).

The horizontal cell contribution to receptive field properties of mouse ganglion cells in the inner retina has remained a controversial topic, including electrical coupling of horizontal cells, which did not seem to play a dominant role in ganglion cell signaling (Dedek et al., 2008). However, a series of recent studies from independent research laboratories provided evidence for horizontal cell signaling as a major influence on ganglion cell receptive field surrounds (Chaya et al., 2017; Drinnenberg et al., 2018; Ströh et al., 2018) with previous inconsistencies in observations likely originating in experimental stimulus properties, such as spatial scales or light levels and the focus on certain ganglion cell types (Ströh et al., 2018). Therefore, the functional consequences of the horizontal cell feature asymmetries observed here would affect the ganglion cell receptive field surround in the upper visual field to be spatially restricted due to smaller cell sizes and lesser extent of electrical coupling (Fig. 6). In the lower visual field, however, the inhibitory feedback signals of horizontal cells distribute across a larger area, causing a much wider surround of postsynaptic cells with potentially lower amplitudes. These ganglion cell receptive field properties have indeed been shown to exist, and they form a gradient across the retina with the most prominent change at the S-opsin transition zone (Gupta et al., 2023), resembling our findings of the horizontal cell gradient and its gap junction coupling asymmetry.

### The cellular basis of the horizontal cell coupling asymmetry

The cellular basis of the difference in electrical coupling is formed by the asymmetric density distribution of the gap junction protein Cx57 at the horizontal cell dendrites. One cannot rule out that some axonal Cx57 has been included in our measurements, but the numbers should be dominated by dendritic gap junctions because of the restricted analysis areas beneath cone pedicles, where most dendritic gap junctions are formed. Interestingly, our observations included a structural change of gap junctions beneath cone pedicles, beyond a mere reduction of Cx57-positive puncta. This was most prominent in the lack of large, elongated ZO-1 plaques. ZO-1 is typically colocalized or closely associated with dendritic horizontal cell gap junctions, where it is thought to form tight junction barriers at the outer perimeter of gap junction plaques at horizontal cell dendrites, but not at their axon terminals (Puller et al., 2009). Thus, both of the most prominent protein components at the horizontal cell dendritic gap junction are regulated to meet the demands of the mouse visual environment. The reduction of ZO-1 in the ventral retina was obvious but less striking than the reduction of Cx57. This is likely resulting from the association of ZO-1 with Cx36-containing gap junctions at OFF bipolar cell dendritic tips beneath the cone pedicle (Puller et al., 2009) combined with an increase in OFF bipolar cell dendritic density in the ventral retina (Camerino et al., 2021; Sharpe et al., 2022). We did not attempt to exclude ZO-1 potentially associated with OFF bipolar cell dendrites from our analysis due to the close spatial vicinity of the two layers of gap junction types.

A compensation of the reduced Cx57 expression by a potential upregulation of another connexin subunit can be ruled out. First, we observed a change in the functional coupling pattern of horizontal cells, which supported the anatomical observations and argued against any compensatory mechanism. Second, Cx50 is another gap junction protein expressed by horizontal cell axons, but it has been shown that it does not compensate for a lack of dendritic Cx57 (Dorgau et al., 2015).

## Conclusions

Mouse horizontal cells are shaping the ganglion cell receptive field surround properties across a spatial gradient of morphological and functional features, which is adapted to the visual scene of the animal. Future studies are required to elucidate the exact interplay of dendritic tree size and potential modulation of electrical coupling, in concert with the complex local and global modes of horizontal cell signaling features (e.g. Jackman et al., 2011; Behrens et al., 2022). In this context, it would be enticing to analyze horizontal cell features in model species from different habitats and life styles, with natural environments which differ from those of the common C57/Bl6 mouse model in terms of scene statistics and general ecology.

## Acknowledgements

We would like to thank Bettina Kewitz and Hannah Käse for excellent technical assistance, Sabrina Duda for expert support with the acquisition and processing of horizontal cell density data, and Asli Pektaş for help with the large-scale quantification of cells in whole-mounted retinas. We also thank Silke Haverkamp and Ben Reese for valuable comments on an earlier version of the manuscript. We acknowledge the Fluorescence Microscopy Service Unit, Carl von Ossietzky University of Oldenburg, for the use of the imaging facilities. Figure 6 was created with BioRender.com. This work was supported by DFG RTG 1885/2 to M.G., U.J.B., K.D. and DFG SFB1372: Magnetoreception and Navigation in Vertebrates, Project 395940726 to K.D. and M.G..

## Author contributions

Conceptualization, U.J.B., K.D., C.P.; Investigation, M.S., A.J.A., L.L.; Formal Analysis, M.S., A.J.A., C.T.B., F.S., Data Curation, C.T.B.; Writing – Original Draft, C.P.; Writing – Review & Editing, all authors; Supervision, M.G., U.J.B., K.D., C.P.

## Supplemental material

### Cx57 antibody validation

Three different antibodies against Cx57 were used in this study (Table 1). They were tested extensively and yielded the same Cx57 staining patterns in the mouse retina as previously published (Hombach et al., 2004; Janssen-Bienhold et al., 2009; Puller et al., 2009). Thus, they were used interchangeably in this study. Two of the polyclonal antibodies against C-terminal peptides of mouse Cx57 were newly raised in guinea pigs (Davids Biotechnologie GmbH, Regensburg, Germany). They were termed CSM and PGS (Fig. S1), and their amino acid sequences are listed in Table 1. The specificity of the antibodies was demonstrated by immunostainings in wild-type and Cx57-deficient mice (Hombach et al., 2004). The common Cx57 immunoreactivity of the two different antibodies was readily observed in wild-type animals following standard protocols (see methods section) but it was absent in retinal sections of Cx57-deficient mice using the same conditions (Fig. S1).

**Figure S1.**
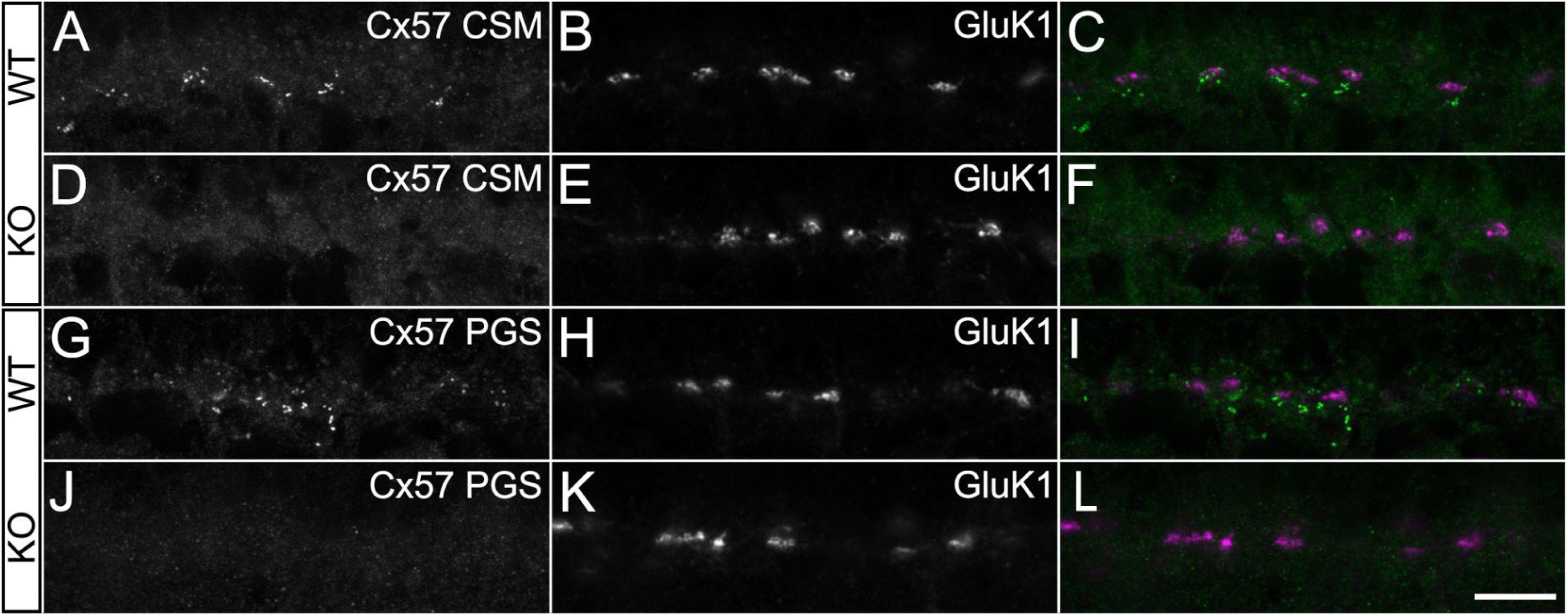
Cx57 antibody validation. **A-F:** Maximum intensity projections of confocal image stacks from vertical cryostat sections labeled with antibodies against the Cx57 CSM epitope and the kainate receptor subunit 1 (GluK1) in retinas from wild-type (WT, A-C) and Cx57-deficient animals (KO, D-F). **G-L:** As in A-F, but with antibodies against the Cx57 PGS epitope. Scale bar: 10 μm.

**Figure S2.**
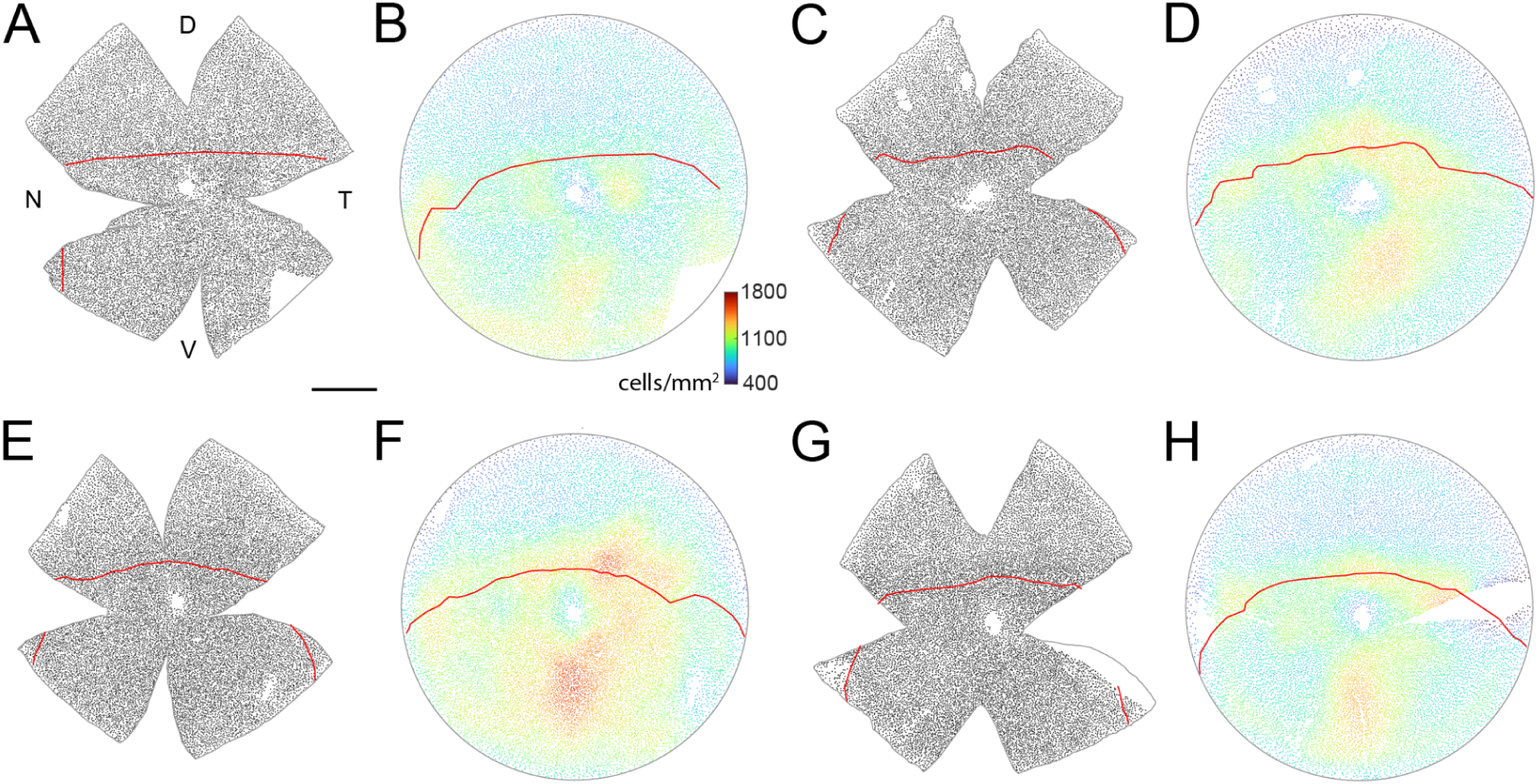
Horizontal cell density distributions. **A:** Left eye of the mouse retina. Each dot represents the cell body position of a horizontal cell marked with CaBP. The red lines in A and B mark the S-opsin transition zone (see also Fig. 1 and 4). To analyze the horizontal cell distribution, all the retinas were oriented as the left eye and added to the average (Fig. 1E), i.e. retinas from the right eye were flipped. D, dorsal; V, ventral; N, nasal; T, temporal. **B**: Azimuthal equal-distant projection of the reconstructed retinal sphere of the data shown in A. The color of the dots represents the local density of the corresponding cell body position. **C–H:** As in A,B but right eye flipped. Scale bar 1 mm in A, applies to A, C, E, G. Color bar in B, applies to B, D, F, H.

**Figure S3.**
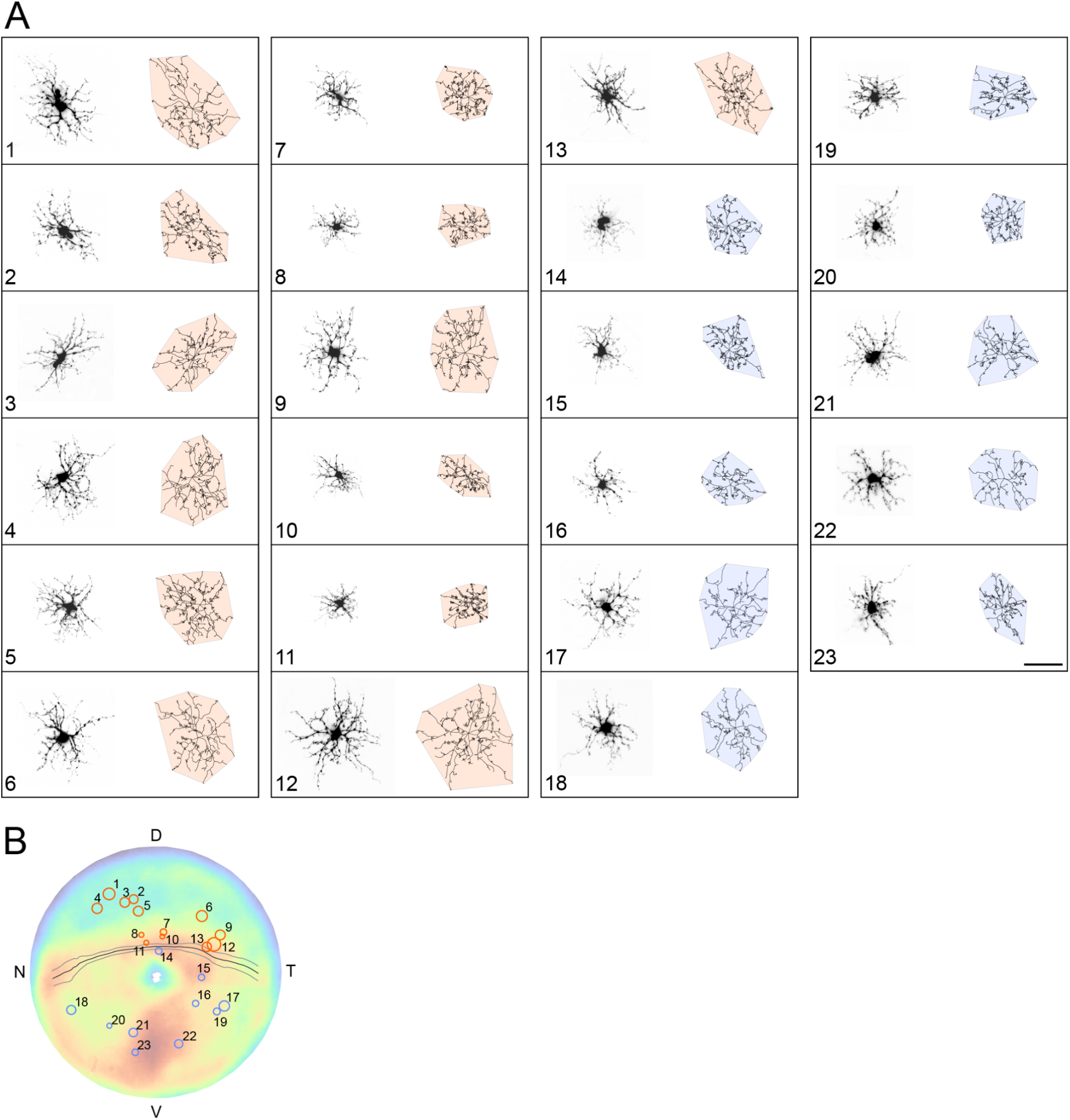
Horizontal cell dendritic tree morphology across the retina. **A:** Maximum intensity projections of image stacks from 23 horizontal cells injected with Alexa dye. Next to a given cell, the skeleton tracing is shown with the corresponding convex hull to calculate the area covered by the dendritic tree. Colors indicate cell position relative to the S-opsin transition zone, either above (orange) or below (blue). **B:** The positions of the cells in A are indicated by numbered circles. The area of a given circle reflects the corresponding dendritic tree area, for better visibility not shown to scale with the averaged density disc from Fig. 1E. Scale bar in A: 50 μm.

